# A pilot study of the impact of smells on the behavior of level 3 autistic children and adolescents during an ongoing task

**DOI:** 10.1101/2023.12.15.571849

**Authors:** M Dantec, F Cazalis, A Peyrical, AC Luisier, C Benabou, S Coeur-Bizot, N Mandairon, M Mantel, M Bensafi

**Affiliations:** Université Claude Bernard Lyon 1, CNRS, INSERM, Centre de Recherche en Neurosciences, de Lyon CRNL U1028 UMR5292, NEUROPOP, F-69500, Bron, France; CAMS UMR8557, CNRS-EHESS, Paris, France; Université de Paris-Nanterre; Association Un pas en avant, Montreuil, France

**Keywords:** Autism, behavior, agitation, screams, smell

## Abstract

The present pilot study aims to perform a first evaluation of the effect of smells on the behavior of autistic children and adolescents (level 3) while they are performing a routine task and are going through a more or less intense crisis situation. To do this, we developed a protocol adapted for a field study in the familiar environment (specialized school) of 8 autistic children and adolescents, where the effect of odors on the behaviors of screaming, agitation, and aggression was assessed. First, it was possible to carry out this protocol on this level 3 autistic population. Second, the results acquired on subjective data collected by the experimenter suggest that odors may reduce certain behavioral traits such as agitation or screaming. These data should be considered as proof-of-concept, with results that need to be confirmed in a larger sample of autistic individuals.

## 1. Introduction

Autism Spectrum Disorder (ASD) is characterized by both (i) deficits in social communication and interaction, and (ii) stereotyped, restricted, and repetitive behaviors, including atypical language and movement, and often atypical sensory issues/behavior (source: DSM-5, American Psychiatric Association, 2022; Ashburner et al, 2013; Kim, 2019). Symptoms that interfere with daily living tasks (e.g., home, school) include stress, anxiety, and emotional dysregulation, which increase the behaviors characteristic of autism, and/or crisis, aggressive behaviors, or agitation (Samson et al, 2015). In addition to these symptoms, one may also cite atypical sensory processing, which includes atypical reaction to smells; a frequent issue in autistic individuals (Kirby et al, 2015).

Given the association between these behaviors and the learning abilities of children and adolescents with autism, it is important to test new strategies to help this population. Moreover, such developments would be particularly relevant in level 3 autistic children and adolescents with communication difficulties, since the self-report approach is problematic in this population compared to the equivalent level 1 and 2 populations which are more widely studied (Russel et al., 2019).

Therefore, the aim of the present pilot study is to develop a methodological approach that will enable an initial assessment of the effect of sensory stimuli such as smells on these behaviors in level 3 autistic individuals. Note that olfaction was chosen because 1/ it is a good sensory input to regulate emotions (Licon et al., 2018a), 2/ children with ASD are sensitive to odors and can experience emotions when smelling odors (Luisier et al., 2015), 3/ odors influence behavior and cognition in children with ASD (Luisier et al., 2019; Parma et al., 2013; Woo & Leon, 2015). To achieve this goal, we conducted a field study in the familiar environment (school) of 8 autistic children and adolescents (level 3), where the effect of the presence of odors on screaming, agitation and aggressive behaviors was assessed. This evaluation of the effect of odors was conducted while the children and adolescents were engaged in a familiar ongoing learning task with an educator.

## 2. Methods

### 2.1. Participants

Eight children and adolescents, seven boys and one girl, with autism spectrum disorders (ASD) were included in this study (age range: 7-17 years; mean+/-SD: 11.88+/-4.01). ASDs were diagnosed by the departmental house for the disabled, and the severity of the disorders was assessed by the ADOS-2 test (Schedule, 2013) (comparison score ranging from 6 to 10 in a scale from 0 to 10, with 10 being the highest rank of autism). RAVEN scores (Raven, 1988) ranged from 16 to 25 (for 3 children the RAVEN score was not interpretable). Finally, the participants’ food neophobia score ranged from 11 to 51 (see Reverdy et al., 2008 for a French version). Note that level 3 autism is difficult to establish on the basis of diagnostic tests alone. This level also corresponds to the level of human assistance required by the autistic person’s condition. The children and teenagers tested had communication difficulties (two were verbal, one could communicate in writing, two could communicate by sign, and three could not communicate by any of these means). When we add that all of them had a severe global handicap, it’s very likely that these participants were at autism level 3. Both parents of the participants gave informed consent to study approved by the Ile de France IV Committee for the Protection of Persons (CPP 2017/55).

### 2.2. Stimuli

During the experimental session, odors were diffused by a diffuser (Aromacare®) remotely controlled by smartphone. The odorants used were cis-3-hexenol (CID 5281167, 1.2% vol/vol; code: CIS) and butanoic acid (CID 264, 0.11% vol/vol; code: BUT). These stimuli were selected on the basis of a previous study (Licon et al., 2018) so that they differed in quality and pleasantness (CIS, smelling like cut grass, pleasant; BUT, smelling rancid butter, cheese, unpleasant).

### 2.3. Experimental design

All participants were exposed to each odor (CIS and BUT), and to a no-odor condition (CON), for a total of three conditions. Each child was exposed to only one condition per day, and the order of presentation of conditions was randomized for each participant. In sum, 24 experimental sessions were conducted (8 children, 3 conditions).

### 2.4. Experimental protocol

The protocol involved three actors: the experimenter (always the same one, first author MD), the educator (the one who use to interact and work with the child), and the child (see Fig 1). In order to familiarize the child with the experimenter, the latter spent time with the child in his or her learning environment in the presence of the educator a few days before the experimental sessions. On the day of an experimental session, the experimenter sat quietly in the room where the educator’s one-on-one lesson with the child was taking place.

**Figure 1.**
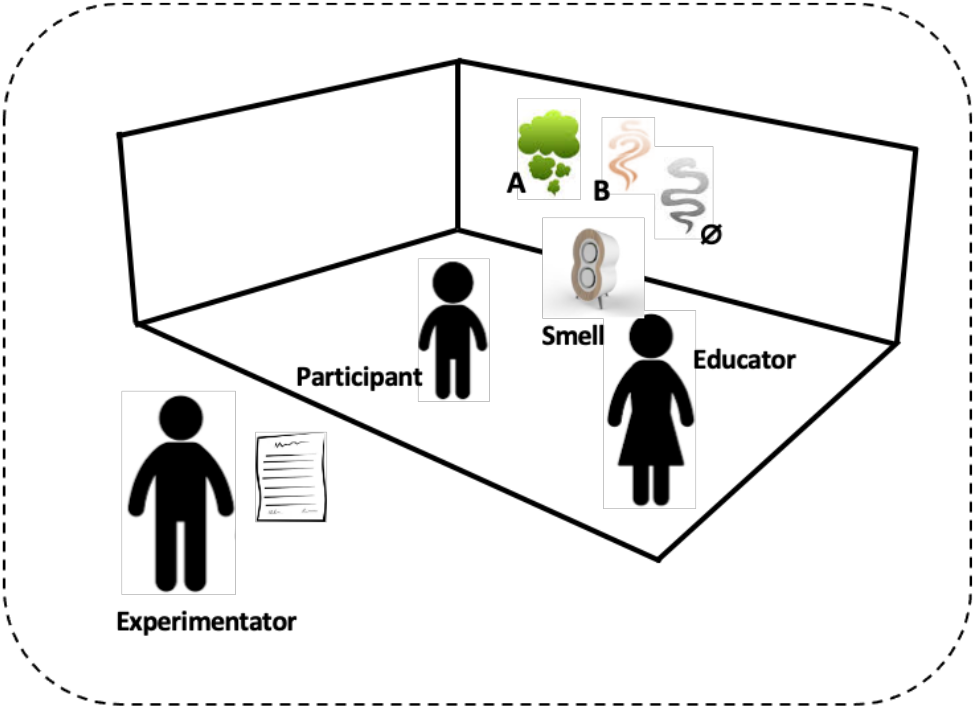
Schema of the experimental protocol.

The educator and experimenter never interacted verbally during the experimental session. When the educator identified that the child was in a crisis situation, the educator made a discreet sign to the experimenter who started the video recording. A camera was positioned in a corner of the room so that the child and the educator were in the camera’s field of view.

From that moment on, the experimental session started. To avoid any olfactory saturation of the room, the experimental session was divided into three periods: a one-minute baseline (before odor diffusion), a one-minute odor diffusion period, and a one-minute post odor diffusion period. The video recordings were cut using these 3 periods: before, during, and after odor diffusion. Although several behaviors were measured (see section 2.5) during these periods, facial expressions could not be measured because the faces had to be blurred for reasons of confidentiality and anonymity.

In addition, during the experimental session, the experimenter evaluated the intensity of the child’s screams, aggressiveness, and agitation during 4 periods: before, during, and 2 min and 5 min after the diffusion of the odor. For this assessment, the experimenter had to use a four-item visual scale: none, low, moderate, high. To perform statistical tests, these qualitative values were transformed into a posteriori numbers (0 for none, 1 for weak, 2 for moderate, and 3 for strong).

In cases of a crisis happening, once the crisis was over, the educator was asked to 1) rate the overall intensity of the crisis (using a qualitative scale: not at all intense, not very intense, intense, very intense), 2) specify the trigger for the crisis (qualitative response), 3) determine whether this type of crisis was very usual, usual, not very usual, or not at all usual, 4) assess whether the child’s behavior was very usual, usual, not very usual, not at all usual during the crisis, and 5) whether it was very easy, easy, difficult, very difficult for the educator to manage this crisis. The educator was also given the opportunity to write free-form remarks.

Finally, at the end of the day, the educator completed a questionnaire about the child’s behavior during the rest of the day (not analyzed here). Items on this questionnaire included: 1/ “Could you tell us how many crisis the child had during the rest of the half-day?”, 2/ “On average, today, were the crises very intense, somewhat intense, not very intense, not at all intense?”, 3/ “Compared to usual, were the crises today more, the same, or less intense?”, 4/ “Did the lunchtime meal go differently from usual? Yes/No (If yes: How was it different?)”, 5/ “Did the child eat all the food offered for lunch? The snack?”, 6/ “On average, today, did the child agree to do the exercises requested: very easily, easily, with difficulty, with great difficulty”, 7/ “Compared to usual, did the child accept more easily, just as easily, or with greater difficulty to do the exercises requested?

### 2.5. Data analysis

The main analyses focused on 2 types of data. First, data related to the experimenter’ assessment of the intensity of screaming, agitation, and aggression were processed using the nonparametric Wilcoxon signed-rank test. Here, for each condition (BUT, CIS, CON) and each variable (screaming, agitation, aggression), the intensity of the behavior during the period before odor diffusion (baseline) was compared with the other 3 periods: during odor diffusion, 2 min after odor diffusion, 5 min after odor diffusion.

Next, behavioral data from the videos were analyzed by extracting a total of 15 parameters from a previous study (Pegoraro et al., 2014). The extraction of the 15 behavioral parameters was performed by 3 judges (co-authors DM, MM, BM) using ELAN software. The analysis was conducted in a blinded fashion with no a priori knowledge of the experimental conditions. The videos were viewed by the 3 judges at the same time, and the target behavior was validated when the agreement of the 3 judges was obtained. Note here that the analyzed behavioral parameters were observed at different frequencies: Echolalia (1.39% of the total 24 sessions and 3 sub-periods before, during and 1 min after odor diffusion), Laughing (1.39%), Rubbing the thighs (4.17%), Tapping the chest / stomach (5.56%), Shaking the head (5.56%), Jumping (6.94%), Autoaggression (8.33%), Flapping (9.72%), Balancing (16.67%), Heteroaggression (16.67%), Striking the head (18.05%), Wiggling the legs (29.17%), Cries (33.33%), Screaming (36.11%) and Automanipulation (47.22%). In order to have a sufficiently powerful statistical analysis, we retained only those variables that appeared in at least 25% of the cases. Moreover, as screaming and crying are both verbalizations, they were combined into a single variable to give more strength to the analysis (by combining the two variables, the frequency of verbalizations reached 50%). Concerning their statistical analysis, the summated durations (in seconds) of the 3 remaining behavioral variables (wiggling the legs, automanipulation, verbalizations) was compared between the period before odor diffusion (baseline) and the other 2 periods (during and after odor diffusion) with a Wilcoxon signed-rank test. This comparison was conducted for each condition (BUT, CIS, CON). Finally, when analyzed, the remaining variables extracted from the different questionnaires were mainly analyzed using descriptive statistics.

## 3. Results

### 3.1. General information

When considering all 24 experimental sessions, children were engaged primarily in school exercise activities (n=21 of 24), or in snack preparation (n=2) or motor/resting activity (n=1). These activities were conducted either in the regular workroom (n=16), in the kitchen (n=1), or in a new room (n=4). The resting activity and the cooking activity were carried out in the motor room and the kitchen respectively. In the 24 experimental sessions, the intensity of the crisis was as follows: not at all intense (25%), weakly intense (37.5%), intense (29.17%), very intense (8.33%). They were caused by several types of inducers: escape or refusal of work (n=10), change of routine or anticipation of future events (n=3), failure to complete a task (n=2), difficulties and concentration related to the task (n=2), other inducers (n=7, e.g. badly cut tape).

### 3.2. Effects of odors

For CIS, agitation was lower 5 min after odor diffusion compared to baseline (before odor presentation) (*W=15, p=0.048*). Three additional trends were observed: decreased aggressiveness during (*W=10, p=0.098*) and 5 min after odor presentation (*W=10, p=0.089*). The other comparisons were not significant (*p>0.05 in all cases*) (Fig 2a.i). The complementary analysis on the behavioral parameters revealed no effect of CIS on wiggling the legs, automanipulation nor verbalizations (*p>0.05 in all cases*) (Fig 2a.ii).

**Figure 2.**
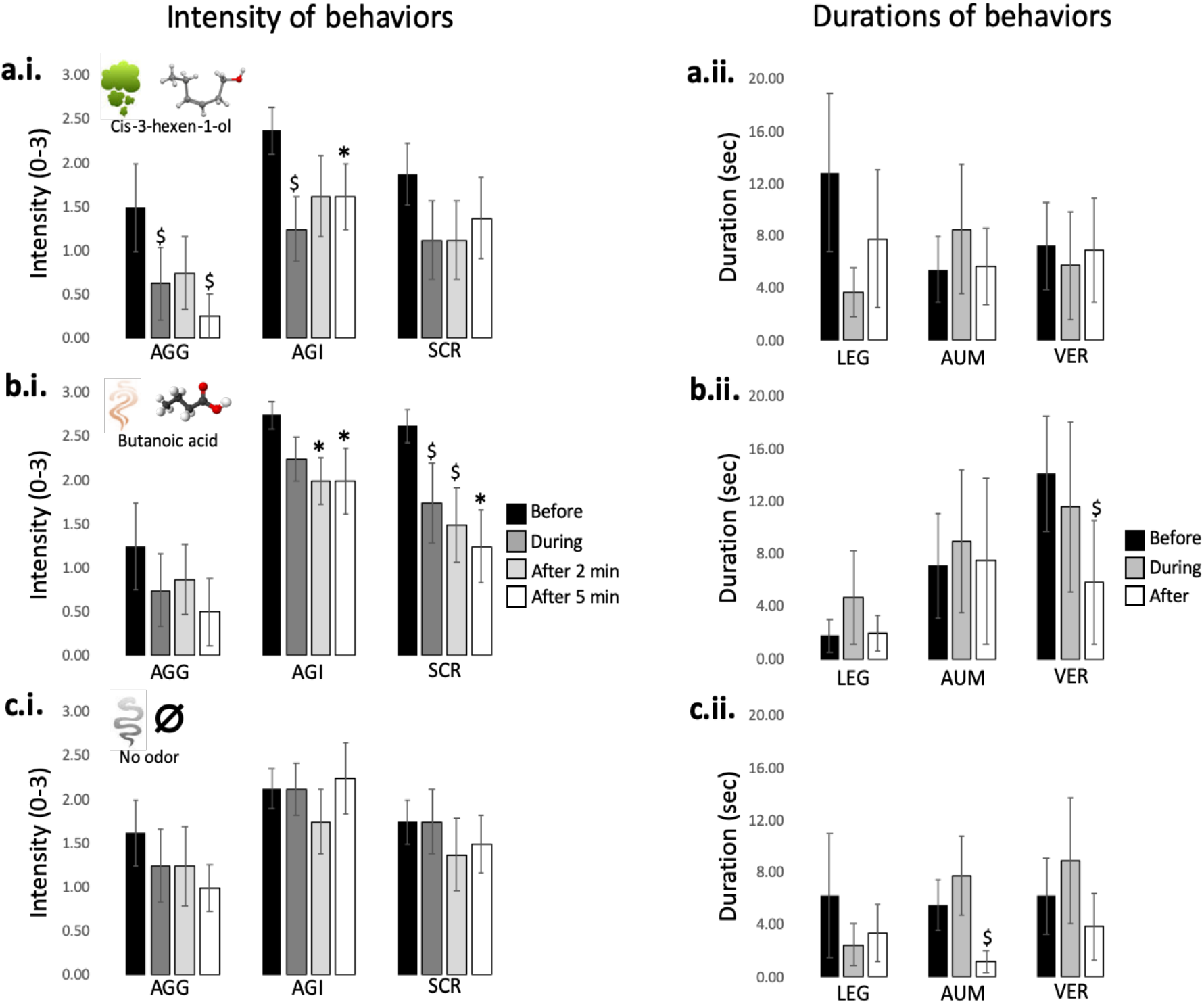
Means and standard errors of intensity of behaviors (“i”) and durations of behaviors (“ii”) for CIS (a), BUT (b) and CON (c). For intensity of behaviors (quoted by the experimenter): AGG means aggressiveness, AGI means agitation and SCR means screaming. For durations of behavior: LEG means wiggling the legs, AUM means automanipulation and VER means verbalizations; *: p<0.05; $: 0.05<p<0.10.

Regarding the BUT condition, compared to baseline (before odor presentation), BUT decreases agitation 2 min (*W=15, p=0.048*) and 5 min (*W=15, p=0.048*) after odor release. Furthermore, screaming was reduced 5 min after odor diffusion compared to baseline (*W=21, p=0.034*). Note that screaming tended to be reduced already during odor presentation (*W=10, p=0.098*) and after 2 min of diffusion (*W=15, p=0.057*). The other comparisons were not significant (*p>0.05 in all cases*) (Fig 2b.i). The analysis of the behavioral parameters showed only one trend: decrease verbalizations when comparing the 1 min post odor diffusion vs. before (*W=15, p=0.059*) (Fig 2b.ii). Finally, for CON, none of the comparisons were significant regarding the intensity of agitation, aggressiveness or screaming (*p>0.05 in all cases*) (Fig 2c.i). For the behavioral parameters, a single trend was observed: decrease automanipulation when comparing the 1 min post odor diffusion vs. before (*W=15, p=0.059*) (Fig 2c.ii).

## 4. Discussion

Since ancient times, humans have been scenting their environment with incense, scented candles, or deodorants to modulate our well-being. Smells can influence our emotions, stress state, and cognitive and social behaviors, even at a low level of consciousness (see for review Licon et al., 2018a).

In this pilot study, we developed a methodological approach that enabled us to conduct a study on the influence of odors on the behavior of children and adolescents with ASD. Overall, the protocol was able to be carried out under experimental conditions involving several constraints constituting a real challenge: i/ the participants had level 3 autism, with severe communication problems, ii/ they were tested in an ecological environment (school, one of their living environments), iii/ they were evaluated in situations of varying degrees of crisis, iv/ the protocol integrated subjective and objective data collection, v/ the diffusion of odors during the protocol. Despite these constraints, the feasibility of the study was not compromised, data could be collected (subjective and objective) and results analyzed and proposed. Whereas these methodological developments provide a good basis for the continuation of similar studies in this level 3 autistic population, the scientific results of our pilot protocol deserve to be discussed.

As our sample size is small our data are very preliminary, nevertheless they suggest that odors can modulate, and perhaps reduce, certain behavioral traits such as agitation or verbalization. These effects were observed mainly on the basis of subjective verbal reports by the experimenter. The data from the behavioral analyses performed by the 3 judges showed few statistical trends, which pointed nevertheless in the same direction as the subjective data collected by the experimenter.

One question raised is the potential mechanisms underlying these effects. It seems that for the autistic children and adolescents tested, ordinary hedonic value is not a key factor in differentiating behavioral changes. CIS and BUT are indeed generally described by neurotypical persons as respectively positive and negative valence odors (Licon et al., 2018b), yet this valence is not reflected in our results. Another possible interpretation would be that odors, regardless of their standard emotional valence, act as attentional distractors and as such, help to regulate the emotional states of children with autism. Theories on emotional regulation support the idea that attentional deployment is a determining factor in regulating our emotions (Gross, 1998). The hypothesis that the odors we used would act as attentional distractors is still speculative at this stage and deserves to be tested using a protocol that includes a larger number of children, with a wider range of autism severity.

The study could be conducted over a longer period of time to assess its replicability and long-term effects on other areas. There are a few scientific studies that have suggested the importance of odors for the social and cognitive development of autistic children. For example, Woo and Leon (2015) exposed children aged 3-12 years with autism to either daily olfactory/tactile stimulation with sensory and cognitive exercises (“enrichment group”) or standard treatment (“control group”). After 6 months of enrichment, the severity of autism (as assessed by the Childhood Autism Rating Scale) was significantly lower in the “enrichment group” than in the “control group”. Another study conducted by Parma and colleagues (2013) showed that in an action task of reaching and grasping an object automatic imitation-an important social skill usually impaired in ASD - could be more efficiently induced in autistic children when the object was associated with the smell of the child’s mother. Those studies and ours all support the idea that smell has to be more systematically thought about when designing individual education programs for autistic children and adolescents.

Regarding the subjective variables analyzed, it is important to raise the issue that the experimenter provided the data on the intensity of the analyzed behaviors. This of course raises the problem of the subjectivity of the measures. To decrease this bias, prior to the start of the tests, the experimenter spent time with each child (at least one session), allowing him/her to become familiar with the children and the rating of aggressiveness, agitation and screaming. However, these observations remain subjective and may be influenced by a range of factors. To circumvent these potential biases, we supplemented this analysis with behavioral measures (summated durations (in seconds) of the 3 behavioral variables namely wiggling the legs, automanipulation, verbalizations). Nevertheless, the small sample size and the inherent difficulties of assessing behaviors by the 3 judges in specific populations such as autistic children only allowed us to observe trends that need to be confirmed.

In conclusion, our study suggests that, from a methodological point of view, hybrid approaches such as ours (combining subjective and objective parameters) can expand the repertoire of methods for studying the behavior, cognition and emotions of level 3 autistic individuals, an under-studied population compared with the same level 1 and 2 populations (Russel et al., 2019). The approach we developed enabled us to assess participants in one of their living environments, under the most ecological conditions possible. The protocol enabled us to test the possibility that odors influence the behavior of children and adolescents with autism while they are engaged in a routine learning task. Preliminary analyses suggested that the presence of odors can modulate the behaviors of children and adolescents engaged in routine tasks, including in situations of intense crisis. Nevertheless, it should be noted that through this article, we provide here only a methodological proof of concept, with scientific results that deserve to be confirmed by a larger evaluation that must consider a larger sample of autistic individuals.

## Acknowledgements

We would like to thank all the children, their parents and educators who made this pilot study possible. The project was funded by the CNRS MITI (AUTON call, EMOTON Project).

## Notes

### Competing Interest Statement

The authors have declared no competing interest.

